# Synaptic Spinules are Reliable Indicators of Excitatory Presynaptic Bouton Size and Strength in CA1 Hippocampus

**DOI:** 10.1101/2022.06.11.495762

**Authors:** Ashley Gore, Amaliya Yurina, Anastasia Yukevich-Mussomeli, Marc Nahmani

**Affiliations:** Division of Sciences & Mathematics, University of Washington | Tacoma, 1900 Commerce St., Tacoma, WA 98402

## Abstract

Synaptic spinules are thin, finger-like projections from one neuron that become fully embedded within the presynaptic or postsynaptic compartments of another neuron. While spinules are a conserved feature of synapses across the animal kingdom, their specific function(s) remain unknown. Recent focused ion beam scanning electron microscopy (FIB-SEM) image volume analyses have demonstrated that spinules are embedded within ∼ 25% of excitatory boutons in mammalian primary visual cortex, yet the diversity of spinule sizes, origins, and ultrastructural relationship to their boutons remained unclear. To begin to uncover the function of synaptic spinules, we sought to determine the abundance, origins, and 3D ultrastructure of spinules embedded within excitatory presynaptic spinule-bearing boutons (SBBs) in mammalian CA1 hippocampus and compare these boutons with presynaptic boutons bereft of spinules (non-SBBs). Accordingly, we performed a comprehensive 3D analysis of every excitatory presynaptic bouton, their embedded synaptic spinules, and postsynaptic densities, within a 5 nm isotropic FIB-SEM image volume from CA1 hippocampus of an adult male rat. Surprisingly, we found that ∼74% of excitatory presynaptic boutons in this volume contained at least one spinule, suggesting that they are fundamental components of excitatory synapses in CA1. In addition, we found that SBBs are 2.5-times larger and have 60% larger PSDs than non-SBBs. Moreover, synaptic spinules within SBBs are clearly differentiated into two groups: small clathrin-coated spinules, and 29-times larger spinules without clathrin. Together, these findings suggest that the presence of a spinule is a marker for stronger and more stable presynaptic boutons in CA1, and that synaptic spinules serve at least two separable and distinct functions.

## Introduction

Synaptic spinules are thin, finger-like invaginating projections from one neuron that become wholly enveloped by the presynaptic or postsynaptic compartments of another neuron (Tarrant and Routtenberg, 1977; Geinisman et al., 1994; Spacek and Harris, 2004). These enveloped protrusions are evolutionarily conserved features across animal phyla, from invertebrate cnidarians (Westfall, 1970; Case et al., 1972) to humans (Zhang and Houser, 1999; Dall’Oglio et al., 2015). Indeed, recent data suggests they are ubiquitous components of mammalian excitatory cortical synapses (Campbell et al., 2020). Yet, synaptic spinule function(s) remain obscure due to a fundamental lack of investigations into their prevalence, morphological characteristics, synaptic specificity, and impact on synaptic function.

A majority of what we know about synaptic spinules *in vivo* comes from data on the protrusions emanating from the dendritic spines of pyramidal cells in the CA1 hippocampus and dentate gyrus of adult rat (Westrum and Blackstad, 1962; Tarrant and Routtenberg, 1977; Geinisman et al., 1994; Spacek and Harris, 2004). Most spinules in these regions seem to project from spines versus other neurites, and nearly all of these spinules are preferentially enveloped by presynaptic boutons (Geinisman et al., 1987; Spacek and Harris, 2004). Furthermore, some of the presynaptic bouton membranes surrounding these spinules are coated in clathrin (Tarrant and Routtenberg, 1977; Sorra et al., 1998; Spacek and Harris, 2004), suggesting an underexplored form of neuronal communication. In addition, the prevalence of spines projecting spinules increases following the induction of long-term potentiation (Schuster et al., 1990; Geinisman et al., 1994; Toni et al., 1999), indicating spinules may participate in activity dependent information transfer and/or circuit remodeling.

However, investigations into spinules within CA1 hippocampus have focused on the spines projecting spinules rather than the prevalence and quantitative morphology of their presynaptic spinule-bearing bouton (SBB) partners. As such, basic questions regarding synaptic spinule structure and function in CA1 remain unexplored, such as: (1) where do spinules enveloped by SBBs originate? (2) what are the morphological features of spinules within SBBs? and (3) how do SBBs and the synapses they make differ from boutons without spinules (non-SBBs)? Here, we investigated these questions through a comprehensive 3D analysis of a 5 nm isotropic focused ion beam scanning electron microscopy (FIB-SEM) open data image volume from the CA1 hippocampus of an adult rat. By analyzing every excitatory presynaptic bouton, postsynaptic density (PSD), and spinule within this volume, we found that SBBs and non-SBBs are quantitatively distinct populations, and that spinules within SBBs come in two discrete ‘flavors’ – small spinules with clathrin-coated tips, and large spinules without clathrin. Taken together, these data suggest that synaptic spinules serve at least two separable and distinct functions, and that the presence of a spinule is a reliable indicator of presynaptic bouton strength and stability within CA1 hippocampus.

## Materials and Methods

### Material

All analyses were performed on a freely available 10.2 × 7.7 × 5.3 *μ*m FIB-SEM image volume (5 nm isotropic voxel resolution) from the *stratum radiatum* of CA1 hippocampus of an adult male rat, originally acquired by Dr. Graham Knott. The protocol that was used for sample fixation, embedding, FIB-SEM preparation, and imaging of this tissue have previously been published (Knott et al., 2011). The full image volume analyzed here is available for download at: https://www.epfl.ch/labs/cvlab/data/data-em/.

### 3D Analyses

Our detailed FIB-SEM analysis protocols have previously been published (Campbell et al., 2020; McCoy and Nahmani, 2021). Briefly, we used ImageJ (i.e., FIJI) to scroll through the image volume at 5 nm z-resolution in order to locate every excitatory synapse and spinule within excitatory boutons in the volume (Schindelin et al., 2012). All 3D analyses were performed and proofread by at least two reviewers to ensure accuracy of traces, the presence/absence of a spinule, and correct spinule origin assignment. Putative excitatory synapses were only included in our analyses if they had (1) parallel alignment of presynaptic and postsynaptic membranes at the active zone, (2) ≥ 3 presynaptic vesicles, (3) a prominent asymmetric postsynaptic density opposite these presynaptic vesicles, and (4) the presynaptic bouton was fully contained (i.e., not cut off) in the image volume. We analyzed each synapse for the identity of the postsynaptic neurite, the presence/absence of a spinule, the presence/absence of a perforated postsynaptic density, and the origin of the spinule (if present). We then assigned a regimented name to each presynaptic bouton based on these characteristics. Spinules were only included in our FIB-SEM analyses if they were observed invaginating into a presynaptic bouton and were then seen in at least one sequential image as a membrane-bound inclusion (i.e., unambiguous visualization of lipid bilayer of spinule surrounded by bouton’s lipid bilayer) within the presynaptic bouton. Spinule origins within FIB-SEM stacks were determined by tracking them within the image volume to their parent neurite or glial process. When it was not possible to identify the parent neurite/glia from which a spinule emerged (i.e., due to an object lacking one or more identifiable morphological criteria by the end of the image volume) we assigned this spinule to the “unknown” origin category. Approximately 7.5% of all SBBs (n = 11) contained a spinule from an “unknown” origin, though most of these SBBs (73%) contained at least one additional spinule from an identified source. PSDs were deemed ‘perforated’ if they displayed one or more separations in the electron-dense region of the PSD, whereas macular PSDs were defined as PSDs bereft of these separations (**Figs. 1B, 2B**, and **3I**). PSD separations needed to be seen in approximately the same x/y location in at least two consecutive serial FIBSEM images (i.e., 10 nm the z dimension) to count as perforations. Every postsynaptic spine was evaluated for the presence of a spine apparatus, identified as tubules of smooth endoplasmic reticulum that needed to be present within both the spine neck and spine head (**Fig. 2C**). To analyze bouton, spinule, and PSD volumes and surface areas, we manually traced and reconstructed 87 SBBs with spinules from AdjAx, AdjS, and PSs parent origins, including their spinules and PSDs, and 51 non-SBBs with their PSDs, using Reconstruct software (Fiala, 2005). 3D reconstruction surfaces were then remeshed and transparency and lighting were adjusted using Blender open-source software (blender.org).

**Figure 1.**
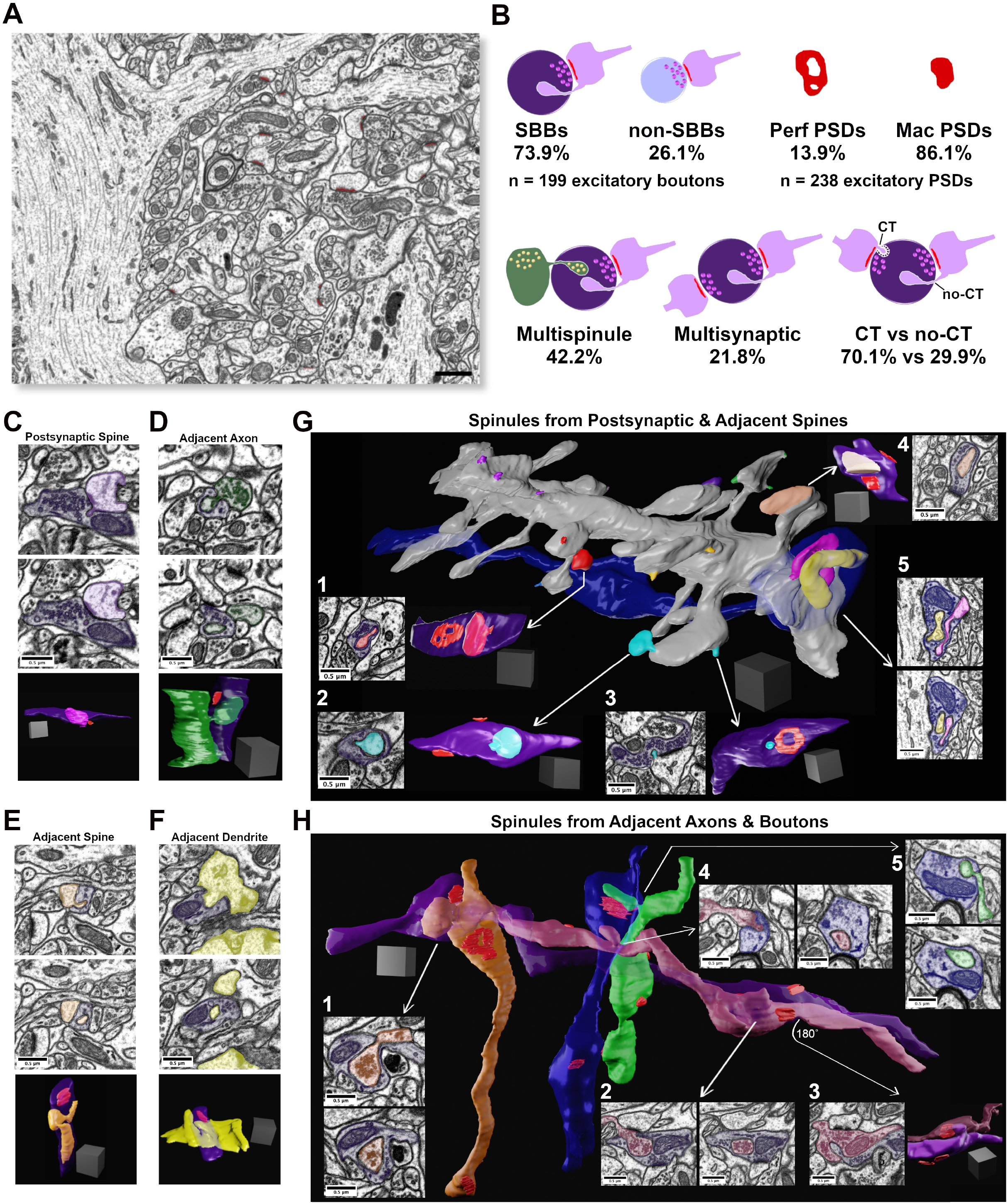
Synaptic spinules are fundamental components of excitatory synapses in CA1. **(A)** Single image from the 5 nm isotropic FIB-SEM image volume of CA1 hippocampus. Excitatory asymmetric postsynaptic densities (PSDs) are pseudocolored red. Note the large apical pyramidal cell dendrites that flank the left and right sides of the image, reducing excitatory synapse density. **(B)** Cartoon showing quantification of the percentage of excitatory presynaptic spinule-bearing boutons (SBBs), presynaptic boutons without spinules (non-SBBs), perforated postsynaptic densities (Perf PSDs), macular PSDs, SBBs enveloping multiple spinules, SBBs with multiple synapses, and spinules with clathrin-coated tips (CT) versus those without clathrin-coats (no-CT). **(C)** Representative example of an SBB (purple) with CT spinules from its postsynaptic spine (PSs; pink) partner. Upper and middle panels are adjacent FIB-SEM images showing the invagination and envelopment of one of three PSs spinules into this SBB. Bottom panel shows the 3D reconstruction of this PSs projecting three spinules into the SBB. Note that this SBB forms a second synapse (PSD in red) with a second spine (not shown). (**D**) Upper and middle panels show adjacent FIB-SEM images of an adjacent bouton (AdjAx, green) projecting a no-CT spinule into an SBB (purple) that has a synapse with a postsynaptic spine (uncolored PSD, top left of SBB). Bottom panel shows the 3D reconstruction of this AdjAx with its large no-CT spinule enveloped by the SBB, and the SBB’s PSD (red) with the postsynaptic spine (not shown). Note the second small CT AdjAx spinule projecting into the very bottom of this SBB. **(E)** Upper and middle panels show two FIB-SEM images of an adjacent spine (no PSD with this SBB; AdjS; orange) projecting (top panel) a large no-CT spinule into an SBB that becomes fully enveloped (lower panel). Bottom panel shows the 3D reconstruction of this AdjS, SBB, and SBB’s PSD (red). Note the spinule’s small (∼30 nm) invagination site (top and bottom panels) that expands into a large, enveloped structure that extends through the majority of this SBB’s length (middle and bottom panels). **(F)** Upper and middle panels show two successive FIB-SEM images of an adjacent dendrite (AdjD, yellow) projecting a no-CT spinule into an SBB (purple), and with the spinule fully enveloped by the SBB (middle panel). Bottom panels show the 3D reconstruction of the surface of this dendrite and its non-synaptic spinule encapsulated by this SBB with a synapse (red) onto a dendritic spine from a separate dendrite (not shown). **(G)** 3D reconstruction of a segment of a dendrite and its spines, a majority of which project spinules into SBBs. All spinules from an individual spine are pseudocolored with the same color. Note that individual spines may project both CT and no-CT spinules (e.g., red and cyan spinules). **(G**_**1**_ **– G**_**2**_) Single FIBSEM sections showing large no-CT spinules within SBBs (purple) emanating from distinct spines (arrows), and the 3D reconstruction of these SBBs (at right) displaying the enveloped spinule and PSDs (red). **(G**_**3**_) FIB-SEM image and 3D reconstruction showing a small CT spinule (cyan) enveloped by an SBB (purple), displaying an annulus-shaped perforated PSD (red). **(G**_**4**_) FIB-SEM image and 3D reconstruction showing an SBB enveloping a spine head spinule (peach) and displaying two macular PSDs (red). (**G**_**5**_) 3D reconstruction (arrow origin) and two successive FIB-SEM images showing an inhibitory presynaptic bouton (blue) with a synapse (symmetric PSD at left of bouton) onto a large pyramidal cell apical dendrite, and two enveloped spinules (pink and yellow) from nearby synaptic spines. **(H)** 3D reconstruction of four excitatory axons (orange, purple, magenta, and green), one inhibitory axon (blue), projecting AdjAx spinules into each other, with visible PSDs (red). Arrows point to FIB-SEM images of the same SBB and AdjAx spinule relationships. **(H**_**1**_) Two successive FIB-SEM images showing an adjacent axon projecting a large no-CT spinule (orange) containing presynaptic vesicles into an SBB (purple). **(H**_**2**_) Two successive FIB-SEM images showing an adjacent presynaptic bouton (magenta) projecting a large no-CT spinule into an SBB (purple). Note that the individual synapses that both boutons have with postsynaptic spines are visible as dark PSDs at top and bottom of image at left. **(H**_**3**_) FIB-SEM image (left) and 3D reconstruction (right) showing the reciprocal relationship to H_2_, with SBB from magenta axon enveloping an AdjAx spinule from the purple axon’s presynaptic bouton and PSDs shown in red. **(H**_**4**_) Two successive FIB-SEM images showing an AdjAx (magenta) with a spinule into an inhibitory presynaptic bouton (blue). **(H**_**5**_) Two successive FIB-SEM images showing another AdjAx (green) with a spinule into this same inhibitory presynaptic bouton (blue) with a synapse (symmetric PSD) onto a dendritic shaft (top FIB-SEM image and red PSD opposite green spinule in 3D reconstruction). Scale bars **(A)** 1 *μ*m, **(C – H)** 0.5 *μ*m; Scale cubes **(C – H)** 0.5 *μ*m per side.

**Figure 2.**
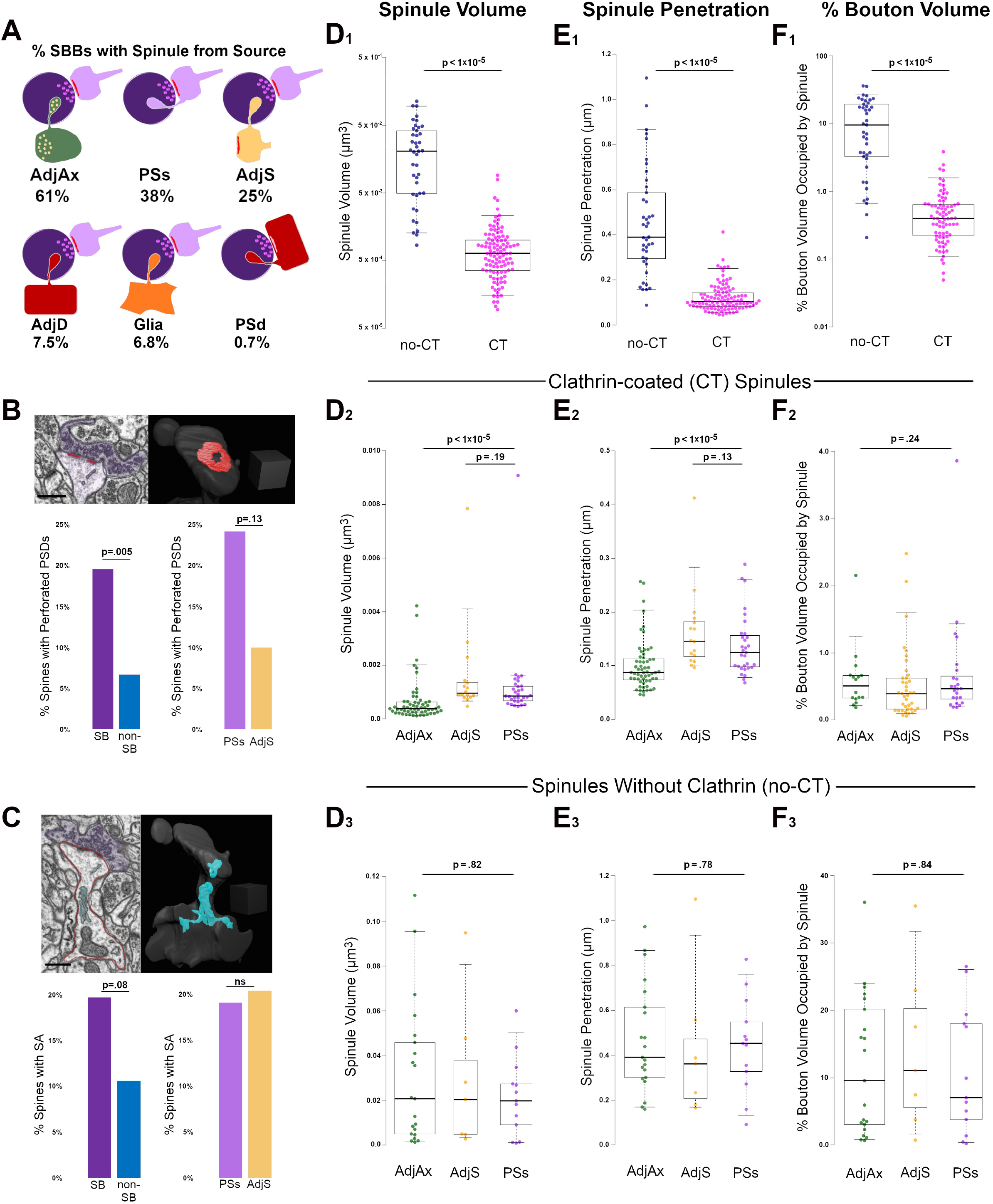
Clathrin-coated (CT) and uncoated spinules (no-CT) are distinct populations within CA1 SBBs. **(A)** Cartoon showing the percentage of SBBs containing at least one spinule from adjacent axons/boutons (AdjAx), postsynaptic spines (PSs), adjacent (no synapse with SBB) spines (AdjS), adjacent dendrites (AdjD), glia, and postsynaptic dendrites (PSd). Percentages reflect the ∼42% of SBBs with spinules from multiple sources. **(B)** Percentage of perforated PSDs in postsynaptic spines in this image volume. Upper left panel shows a FIB-SEM image of a postsynaptic spine (pink) with a perforated PSD (red) and its presynaptic SBB partner (purple). Note the CT spinule projecting just to the right of the PSD. Upper right panel shows 3D reconstruction of spine and PSD from FIB-SEM image; note this is the same spine as shown in Fig.1G_2_ – G_3_. Lower left panel shows the percentages of spinule-bearing (SB) and non spinule-bearing (non-SB) postsynaptic spines displaying perforated PSDs. Lower right panels show the percentages of spines projecting PSs spinules and AdjS spinules into SBBs that display perforated PSDs. **(C)** Percentage of spines with a spine apparatus in this image volume. Upper left shows a FIB-SEM image of the same postsynaptic spine (outlined in pink) as in (B), displaying a spine apparatus (cyan), with a synapse onto an SBB (purple). Upper right panel shows the 3D reconstruction of this spine and its spine apparatus. Lower left panel shows the percentages of SB and non-SB spines displaying a spine apparatus, while lower right panels show the percentages of spines projecting PSs and AdjS spinules containing a spine apparatus. **(D)** Spinule volume comparisons between no-CT (blue) and CT spinules (pink) **(D**_**1**_), between CT spinules of AdjAx (green), AdjS (yellow), and PSs origins (magenta) **(D**_**2**_), and between no-CT spinules of AdjAx, AdjS, and PSs origins **(D**_**3**_). For all boxplots, center lines show medians, box limits indicate the 25^th^ and 75^th^ percentiles, whiskers extend to the 5^th^ and 95^th^ percentiles, width of boxes is proportional to the square root of the sample size, and raw data points are plotted as filled circles. **(E)** Comparisons of spinule penetration distance into their SBBs between no-CT and CT spinules **(E**_**1**_), between CT spinules of AdjAx, AdjS, and PSs origins **(E**_**2**_), and between no-CT spinules of AdjAx, AdjS, and PSs origins **(E**_**3**_**). (F)** Comparisons of the percentage of SBB volume occupied by no-CT and CT spinules **(F**_**1**_), CT spinules of AdjAx, AdjS, and PSs origins **(F**_**2**_), and no-CT spinules of AdjAx, AdjS, and PSs origins **(F**_**3**_). Scale bars **(B & C)** 0.5 *μ*m; Scale cubes **(B & C)** 0.5 *μ*m per side.

### Statistics

Statistical tests, means ± standard error of the mean (SEM), p-values, effect sizes, and statistical power for all comparisons are listed in **Table 1**. For all tests, α was set at 0.05, except where Bonferroni or Yates corrected p-values were used as appropriate and listed in Table 1.

**Table 1.**
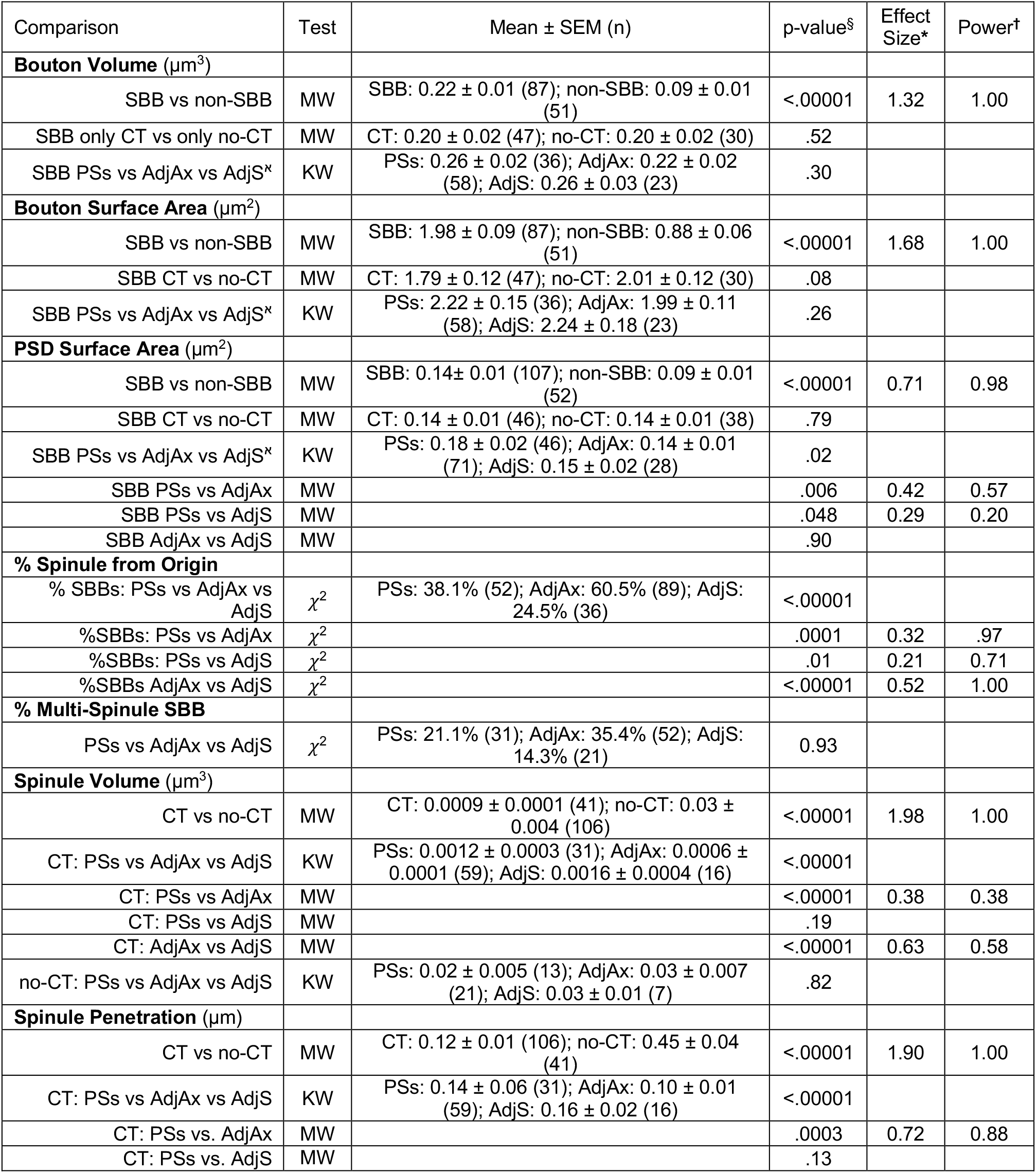
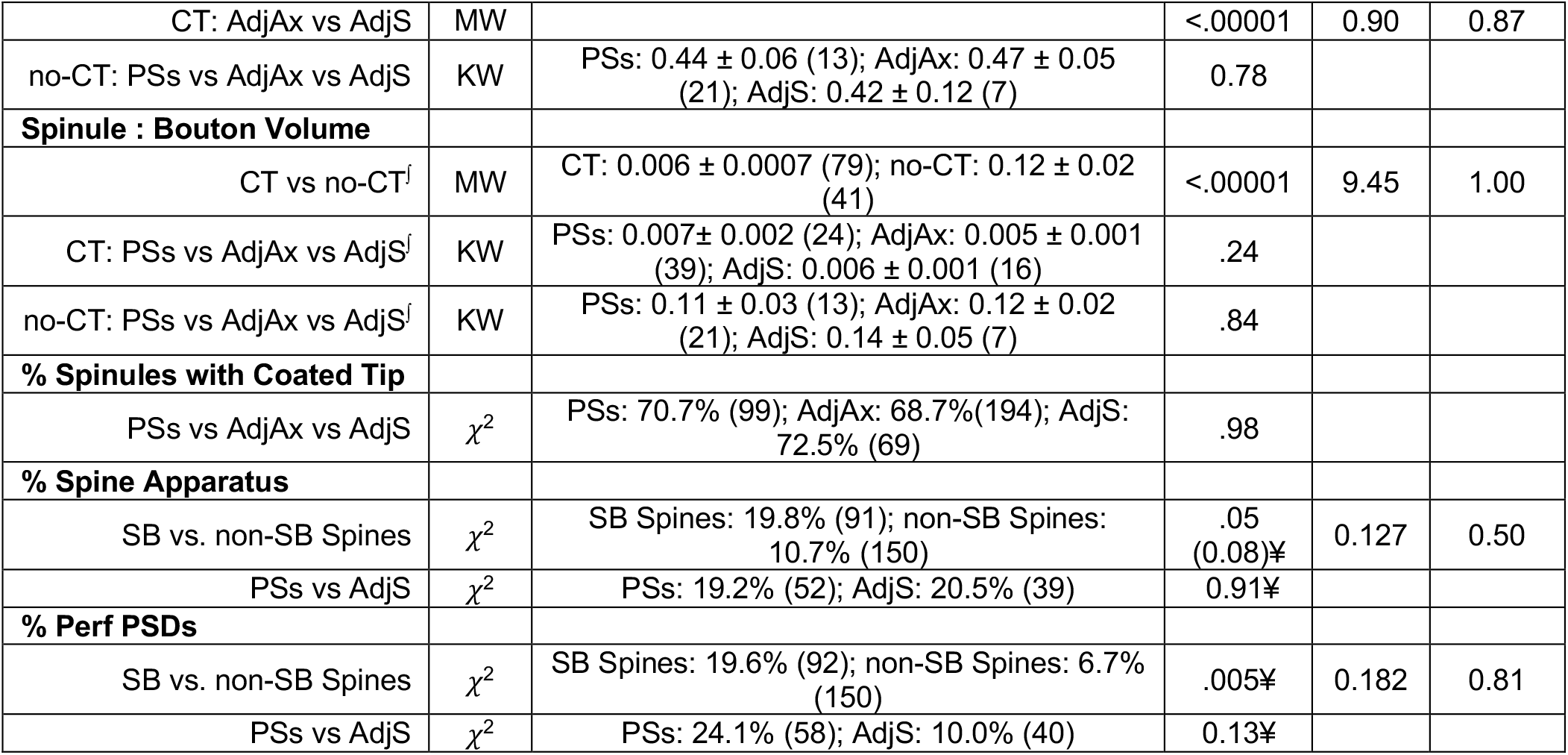
Means and Descriptive Statistics. ^§^*α* set at 0.05, Bonferroni corrected value = 0.025 and .017 for two and three group comparisons, respectively. ^**†**^Post hoc achieved power at α = 0.05, two-tailed; **\***Hedges’ g for MW, *w* for *χ*^2^; MW = Mann-Whitney U test; KW = Kruskal-Wallis test; *χ*^2^ for Chi-Square test. ¥ Yates corrected p-value; ^**ℵ**^Some SBBs are counted multiple times in this comparison because they contain spinules from multiple origins. ^∫^If multiple spinules of the same type (e.g., CT) were present within the same bouton, their volumes were summed for this comparison. SBB = Spinule-Bearing Bouton; CT = Clathrin-coated spinule; SB = spinule-bearing; PSs = postsynaptic spine; AdjAx = adjacent axon; AdjS = adjacent spine.

## Results

### Synaptic Spinules Are Fundamental Components of Excitatory Synapses in CA1

To investigate the origins, ultrastructural diversity, and prevalence of spinules with hippocampal excitatory SBBs, we took advantage of a freely available 10.2 × 7.7 × 5.3 *μ*m FIB-SEM image volume with 5 nm isotropic voxel resolution, from *stratum radiatum* of CA1 hippocampus of an adult male rat (**Fig. 1A**). To determine the prevalence of spinules within the excitatory boutons in this data set, we performed a 3D analysis to locate every excitatory presynaptic bouton within this image volume that contained a fully enveloped spinule (see Methods). Surprisingly, we found that of the total excitatory presynaptic boutons (n=199) in this CA1 image volume, a large majority (∼74%) contained at least one spinule, while a minority lacked spinules (∼26%; **Fig. 1B**). By tracing each spinule back to the neurite it emanated from within the 3D image volume, we determined that spinules embedded within these SBBs originated from a diverse array of sources, including: spinules projected from postsynaptic spines (PSs, **Fig. 1C**), adjacent axons / boutons (AdjAx, **Fig. 1D**), adjacent (i.e., no synapse with SBB) spines (AdjS, **Fig. 1E**), adjacent dendritic shafts (AdjD, **Fig. 1F**), postsynaptic dendrites (PSd), glia, and a small percentage of indeterminate origin (7.5%; see Methods). Yet, SBBs predominantly contained spinules from just three sources, AdjAx (61%), PSs (38%), and AdjS (25%), with ≤ 7.5% of SBBs containing a spinule from any other single source (**Fig. 2A**, Table 1). Note that these percentages reflect the ∼42% of SBBs that contained more than one spinule (mean = 1.7, range = 1 – 6, spinules per SBB). Interestingly, AdjAx spinules often contained presynaptic vesicles and/or a smooth endoplasmic reticulum within their lumens, a possible route for communication between boutons (**Fig. 1D, H**_**1**_ – **H**_**2**_). Furthermore, while our analyses were primarily focused on excitatory synapses, we observed inhibitory presynaptic boutons with enveloped PSs and AdjAx spinules from excitatory neurons (**Fig. 1G**_**5**_, **H**_**4**_, and **H**_**5**_), demonstrating the potential for novel forms of excitatory to inhibitory neuron communication. These data suggest that spinules are fundamental components of excitatory synapses in CA1 hippocampus, and that most spinules within SBBs come from dendritic spines, presynaptic boutons, and axons.

During our analysis, it became apparent that spinules within SBBs often displayed clathrin-coats along the leading edge of the invaginating presynaptic membrane (**Fig. 1B, 1C, 1G**_**3**_), as previously observed in rat CA1 (Spacek and Harris, 2004). These clathrin-tipped (CT) spinules appeared relatively small, versus spinules without clathrin (no-CT), and CT spinules emanated from every identified source of spinules except postsynaptic dendritic shafts. There were no detectable differences in the percentages of CT spinules from the three main spinule sources (p = .98, *χ*^2^ = 0.04; 68.7%, 72.5%, 70.7%, for AdjAx, PSs, and AdjS spinules, respectively). We quantified the percentage of SBBs (n = 147) containing at least one CT spinule versus those only containing no-CT spinules and found that a majority of SBBs contained CT spinules (70.1 vs. 29.9 %, CT vs. no-CT, respectively, n = 302 spinules from all sources; **Fig. 1B**). In addition, we found that while most presynaptic boutons had postsynaptic partners whose PSDs (n = 238 total PSDs) displayed a macular morphology (∼86%), some of the PSDs associated with these excitatory presynaptic boutons displayed perforations (∼14%, **Fig. 1B**), similar to a previous 3D analysis of excitatory PSDs in adult rat CA1 (8 – 12%; (Sorra et al., 1998). Given these differences in the percentages of SBBs containing spinules from distinct origins, and the presence or absence of a clathrin coat, we sought to determine if there were quantitative differences between spinules from different origins, or between spinules with and without a clathrin coat, that might hint at their function. Accordingly, we restricted the rest of our analyses to SBBs containing spinules from the three major neurite origins (AdjAx, PSs, and AdjS) to achieve reasonable statistical power across our comparisons (see Methods and Table 1).

### Clathrin-coated and Uncoated Spinules are Distinct Populations

To determine whether spinules from different origins or those with clathrin coating are morphologically distinct, we three-dimensionally reconstructed 87 SBBs containing 147 AdjAx, PSs, and AdjS spinules. All statistical tests, sample sizes (n), p-values, effect size, and power for these and all further comparisons can be found in **Table 1**. Our analyses revealed that no-CT spinules were 29-times larger, projected 3.8-times farther into their SBB, and occupied 21-times more volume than CT spinules (p < 1×10^−5^; **Fig. 2D**_**1**_, **E**_**1**_, and **F**_**1**_), indicating that CT and no-CT spinules are separate populations. We hypothesized that if CT and no-CT spinules that emanate from different origins have distinct functions (e.g., large spinules that enhance synaptic stability vs. small spinules designed for transendocytosis), they would likely display distinct morphologies. Thus, we compared CT spinules and no-CT spinules across origin categories (i.e., AdjAx, PSs, and AdjS). We found that CT spinules from AdjAx are ∼2-times smaller than CT spinules from PSs and AdjS (p < 1×10^−5^; **Fig. 2D**_**2**_, **E2**, and **F2**), suggesting CT spinules from boutons and axons may have separable functions from CT spinules originating from spines.

However, AdjAx, PSs, and AdjS CT spinules occupied a similar percentage of their SBBs, indicating that AdjAx CT spinules may preferentially target SBBs that are smaller than those containing PSs and AdjS spinules. Although no-CT spinules were larger than their CT spinule counterparts, no-CT spinules from AdjAx, AdjS, and PSs origins were indistinguishable from each other in volume, bouton penetration, and the percent volume they occupied within their SBBs (p = .80; **Fig. 2D**_**3**_, **E**_**3**_, and **F**_**3**_). Together, these results argue for two separable and distinct forms of spinules within the excitatory SBBs in CA1: small spinules participating in clathrin-mediated transendocytosis and large spinules devoid of clathrin that invaginate deep (∼450 nm) into their enveloping SBBs.

### Spinule-Bearing Spines Associated with Perforated PSDs and Spine Apparatuses

Perforated PSDs are associated with increases in cortical activity (Calverley and Jones, 1990; Geinisman et al., 1993; Toni et al., 1999), and recent findings have suggested that spinule-bearing spines have higher rates of perforated PSDs (Campbell et al., 2020; Zaccard et al., 2020). Thus, we next sought to examine whether PSs and AdjS spinule-bearing spines (SB spines, n = 92) differed from spines without spinules (non-SB spines, n = 150) on this correlative measure of heightened activity state. We found that while only 6.7% of non-SB spines had perforated PSDs, 19.6% of SB spines had perforated PSDs (p = .005). Interestingly, spines with PSs spinules showed a trend toward higher rates of perforated PSDs versus spines projecting AdjS spinules (24.1% vs. 10.0%, p = .13, n = 58 and 48, for PSs vs. AdjS, respectively; **Fig. 2B**). However, given our relatively small sample size, this latter result awaits confirmation from a larger dataset (see Discussion). In sum, these findings intimate that spinules emanating from spines play a role in the dynamic partitioning of the PSD.

The presence of a spine apparatus is also associated with large spines that have recently experienced high levels of synaptic activity (Chirillo et al., 2019; Perez-Alvarez et al., 2020). Accordingly, we quantified the percentages of SB and non-SB spines containing a spine apparatus. We found that 19.8% of SB spines had a spine apparatus, while only 10.7% of non-SB spines had a spine apparatus (p = 0.08; n = 91 and 150, SB and non-SB spines, respectively, **Fig. 2C**). PSs and AdjS projecting spines had similar percentages of spine apparatuses (19.2% vs. 20.5%, p = .91). Thus, SB spines showed a trend toward displaying an organelle associated with robust levels of synaptic activity at higher percentages than non-SB spines.

### Synaptic Spinules are Markers of Stronger Synapses

Our findings on spinule populations in CA1 highlighted the diversity of spinule morphologies, associated postsynaptic organelles, and their potential functions, prompting us to ask whether select presynaptic boutons (e.g., SBBs containing large no-CT spinules) might be larger than SBBs with small spinules or non-SBBs. Thus, we measured the volume and surface areas of SBBs containing AdjAx, AdjS, and PSs spinules (n = 87) and presynaptic non-SBBs (n = 51; **Fig. 3A** - **3F**). We found that SBBs had 2.5-times larger volumes and 2.3-times larger surface areas than their non-SBB counterparts (p < 1×10^−5^; **Fig. 3G and 3H**). However, we did not detect an influence of spinule origin on SBB size (p = .30 and .26, for PSs vs AdjAx vs. AdjS volume and surface area, respectively, **Table 1**). We next compared the PSD surface areas of SBBs versus non-SBBs as a measure of their respective synaptic strengths (Branco et al., 2010; Holderith et al., 2012; Araya et al., 2014; Meyer et al., 2014; Holler et al., 2021). Similar to SBB volume and surface area, we found that PSD surface areas at SBB synapses were 1.6-times larger than PSD surface areas at non-SBB synapses (p < 1×10^−5^, n = 107 and 52 for SBB and non-SBB PSDs, respectively; **Fig. 3I and 3J**). Hence within rat CA1, the presence of a spinule within an excitatory presynaptic bouton is a marker for a larger and stronger synaptic contact.

**Figure 3.**
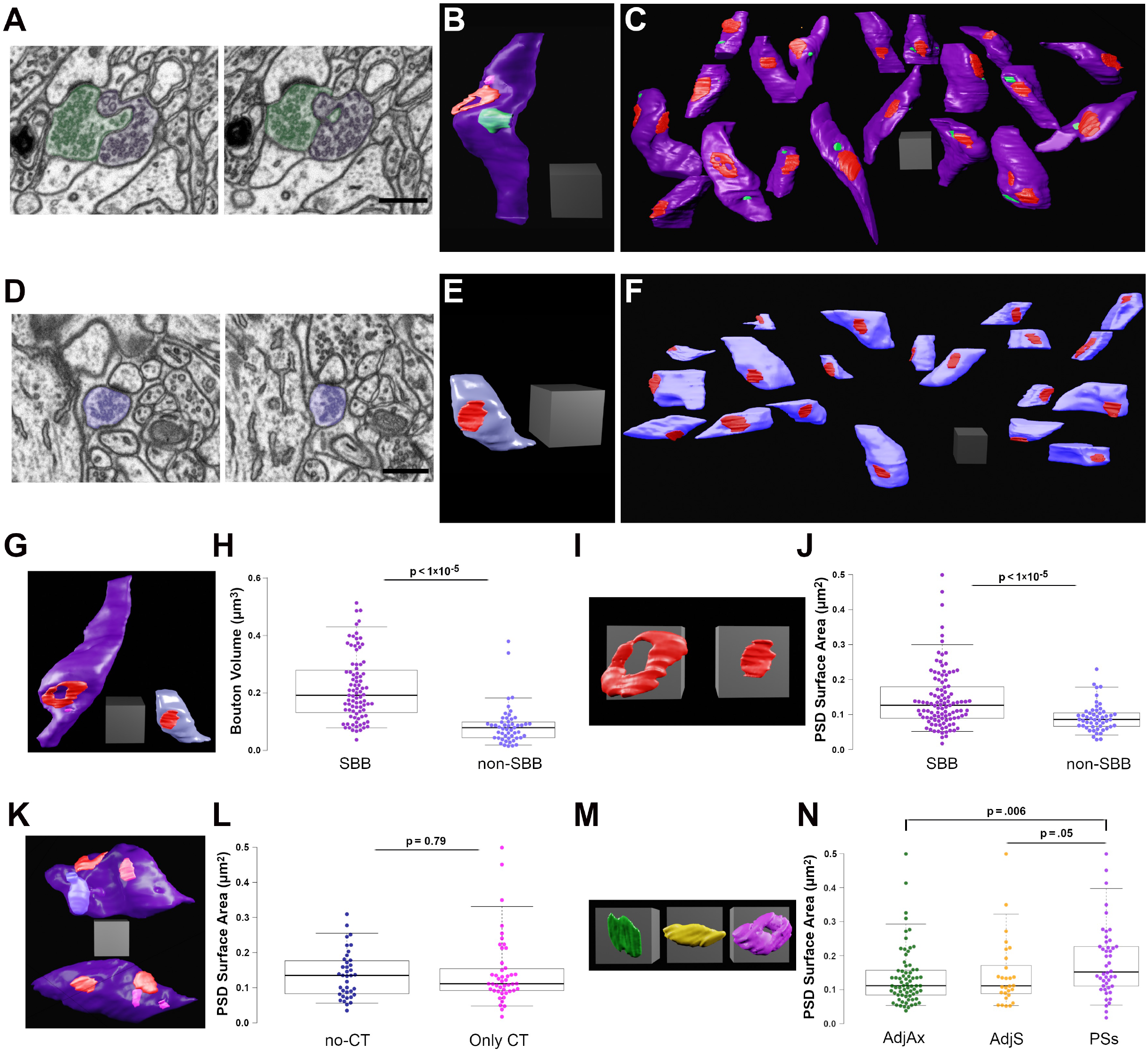
Spinule-bearing presynaptic boutons (SBBs) are larger and stronger than presynaptic boutons without spinules (non-SBBs). **(A)** Successive FIB-SEM images showing a spinule from an adjacent presynaptic bouton (AdjAx; green) with a spinule projecting into and enveloped by an SBB (purple). **(B)** 3D reconstruction of the same SBB (purple) and AdjAx spinule (green) as shown in (A), displaying an annulus-shaped perforated PSD (red) and a CT spinule (pink) from a postsynaptic spine (not shown). **(C)** Field of 19 3D reconstructed SBBs (purple) in their approximate neuropil locations within the FIB-SEM image stack; some SBBs have been rotated to display their PSDs (red) and spinule invagination sites (green). **(D)** Successive FIB-SEM images showing a presynaptic non-SBB (light blue) with a synapse (dark PSD above bouton) onto a postsynaptic spine. Note that this postsynaptic spine is enveloped by an inhibitory presynaptic bouton (top of right FIB-SEM image) and has a long no-CT spinule into this inhibitory bouton (see yellow spinule in Fig. 1G_5_). **(E)** 3D reconstruction of the same non-SBB (light blue) and PSD (red) as shown in (D). **(F)** Field of 20 non-SBBs (light blue) in their approximate neuropil locations within the FIB-SEM image stack; some SBBs have been rotated to display their PSDs (red). **(G)** Same 3D reconstructions of SBB and non-SBB shown in (B) and (E), shown side by side for comparison. (**H)** Box plots showing the presynaptic bouton volume for SBBs and non-SBBs (n = 87 and 51, respectively). For all boxplots, center lines show medians, box limits indicate the 25^th^ and 75^th^ percentiles, whiskers extend to the 5^th^ and 95^th^ percentiles, width of boxes is proportional to the square root of the sample size, and raw data points are plotted as filled circles. **(I)** Zoomed-in view of the 3D reconstructions in (G) showing PSDs from an SBB (left) and a non-SBB (right). **(J)** Boxplots of PSD surface area for SBBs and non-SBBs (n = 107 and 52, respectively). **(K)** 3D reconstructions of SBBs with no-CT (above, light blue) and CT (below, pink) spinules, displaying their PSDs (red). **(L)** Boxplots of PSD surface area for SBBs with only no-CT spinules and SBBs with only CT spinules. **(M)** 3D reconstructions of representative PSDs from SBBs containing AdjAx (green), adjacent spine (AdjS, yellow), and postsynaptic spine (PSs, magenta) spinules. **(N)** Boxplots of PSD surface area for SBBs containing spinules from AdjAx, AdjS, and PSs origins. Scale bars **(A & D)** 0.5 *μ*m; All scale cubes = 0.5 *μ*m per side.

Since we had determined that no-CT spinules were 29-times larger than CT spinules, we hypothesized that SBBs enveloping no-CT spinules might be larger than those containing CT spinules. Yet surprisingly, in comparing SBBs that only contained small CT spinules (n = 47) with those that only contained larger no-CT spinules (n = 30), there were no detectable differences in their volumes (p = .52) or their PSD surface areas (p = 0.79; **Fig. 3K** and **3L**). However, we found a small but intriguing trend for no-CT containing SBBs possessing larger surface areas than SBBs with CT spinules (p = .08, 2.01 ± 0.12 vs. 1.79 ± 0.12, for no-CT and CT, respectively), intimating that large spinules might impact the morphological complexity of their SBBs. In addition, in comparing the PSD surface areas from SBBs containing spinules from AdjAx, AdjS, and PSs sources, we found that PSDs from SBBs with PSs spinules were 33% larger than PSDs from SBBs containing AdjAx spinules and showed a trend toward having larger PSDs than SBBs containing AdjS spinules (p = .006 and .05 for PSs vs. AdjAx and AdjS, respectively, **Fig. 3M** and **3N**). Together, these data suggest that spinule presence, regardless of size, is predictive of a distinct class of larger and stronger presynaptic boutons, and that SBBs with PSs spinules may possess the largest PSDs within the CA1 SBB population.

## Discussion

In this study, we sought to answer three questions: (1) Where do the spinules within excitatory SBBs in CA1 originate? (2) Do spinules within these SBBs display distinct morphologies? (3) How do SBBs and their PSDs differ from non-SBBs? Our 3D analyses of all the excitatory synapses within this 419 *μ*m^3^ 5 nm isotropic CA1 FIB-SEM image volume revealed that spinules within SBBs predominantly originate from three sources: adjacent excitatory axons / boutons, postsynaptic dendritic spines, and adjacent (non-synaptic) spines. In addition, we found that spinules come in two main categories: small clathrin-coated projections that are likely involved in transendocytosis (Spacek and Harris, 2004), and 29-times larger uncoated spinules that project deep into their host SBBs. Furthermore, we show that SBBs are 2.5-times larger and have 60% larger PSDs than their non-SBB counterparts within the same image volume.

Together, these findings demonstrate that the presence of an encapsulated spinule is a robust predictor of excitatory presynaptic bouton size and strength, differentiating large and strong SBBs from smaller, weaker, and potentially less stable non-SBBs. Moreover, these data intimate that spinules serve at least two functions: small spinules involved in macromolecular communication through clathrin-mediated vesicular transendocytosis, and large spinules that significantly increase neuron-to-neuron extrasynaptic membrane interface to potentially enhance connection stability and/or allow for a unique form(s) of neuronal communication.

### Experimental Limitations

These 3D analyses were conducted on one image volume from the CA1 hippocampus of an adult male rat. We chose this approach because it allowed us to manually analyze every excitatory bouton and synaptic spinule within an image volume of CA1 within reasonable time constraints. We concluded that a manual analysis approach in an image volume with high spatial resolution (i.e., < 15 nm voxels) was best suited to our questions because spinule invagination sites into SBBs are often less than < 40 nm wide (e.g., Fig. 1C, 1D, 1E, and 1H_5_). However, while we may have captured the variability inherent within this animal’s excitatory synapses in CA1, we could not capture inter-animal variability given that these data were from a 419 *μ*m^3^ volume from a single animal. For example, this image volume will likely have a lower excitatory synapse density than some other volumes of CA1 due to the large apical pyramidal dendrites that flank the majority of its images (Fig. 1A), at times reducing the area of synaptic neuropil by > 50%. However, the averages and ranges for the excitatory bouton volumes and PSD surface areas we analyzed in this volume are consistent with previous 3D ultrastructural analyses of excitatory presynaptic boutons in rat CA1 (Shepherd and Harris, 1998; Grillo et al., 2018), indicating that the synaptic ultrastructure within our image volume is comparable with that of other analyses of the excitatory synapse population in rat CA1.

Our analyses revealed large differences in bouton volume and surface area between SBBs and non-SBBs, SBB and non-SBB PSDs, and CT vs. no-CT spinules (**Table 1**). Yet, as we expanded our analyses to include combinations of multiple variables/characteristics (e.g., perforated synapses from AdjS containing SBBs), our sample size was necessarily restricted, effectively decreasing our statistical power for two noted comparisons: the percentage of spine apparatuses within SB and non-SB spines, and the percentage of perforated PSDs from PSs and AdjS spines. Since the overall percentage of spines that contain a spine apparatus or perforated PSD in CA1 is small (7-20% in this study), it is difficult to achieve high levels of statistical power when comparing subgroups of spines (e.g. AdjS projecting spines) even when the computed difference between groups is large (e.g., 2-fold increase). For example, to achieve a statistical power of 0.8 for our spine apparatus comparisons, given the 2-fold difference between groups we observed (i.e., *χ*^2^ effect size = 1.3 - 1.4), would require a doubling of the total area used in this analysis. As such, these two comparisons (% spine apparatus between SB and non-SB spines, and % perforated PSDs between PSs and AdjS spines), while intriguing, await confirmation or refutation from future analyses of large volume high resolution (< 15 nm voxel) FIB-SEM datasets.

### SBBs and the Potential for Axo-axonic Communication

In our previous 3D analysis of SBBs within a FIBSEM image volume of late adolescent ferret primary visual cortex, we found that the primary sources of spinules within SBBs were from AdjAx (35.2%) and PSs (37.0%), AdjD (14.8%), and AdjS (7.4%) (Campbell et al., 2020). Thus, while SBBs in each brain area may preferentially envelop small percentages of spinules from specific sources based on distinct microcircuit functions, AdjAx and PSs spinules seem to be primary sources for spinules within SBBs across brain areas. Indeed, AdjAx spinules are the primary origin for spinules within SBBs across the postnatal development of ferret primary visual cortex (Campbell et al., 2020). Interestingly, it has been reported that only ∼12% of spinules emerge from axons and presynaptic boutons in adult rat CA1, with ∼86% projected from dendritic spines (Spacek and Harris, 2004). Here, we found that 61% of spinules within SBBs in adult CA1 originated from adjacent axons and presynaptic boutons. This discrepancy in the percentage of spinules emerging from adjacent axons and those enveloped by SBBs could be due to our ability (with 5 nm isotropic voxel resolution) to detect spinules with invagination widths that were smaller than the section thickness (50 nm) used in the only other systematic investigation of spinules origins in CA1 (Spacek and Harris, 2004). Regardless, data from our analyses of five FIB-SEM image volumes demonstrate that AdjAx spinules are a primary source of spinules within excitatory SBBs in adult rat CA1 and developing ferret primary visual cortex (Campbell et al., 2020).

Indeed, these findings raise the possibility that axo-axonic communication may be more common than presently understood. While there are examples of axo-axonic communication across brain regions (Cover and Mathur, 2021), axo-axonic synaptic contacts likely comprise a small minority of the total excitatory and inhibitory synapses across the brain (Somogyi et al., 1982; DeFelipe and Fariñas, 1992; Inan et al., 2013). Thus, if as we’ve shown 45% of CA1 excitatory synapses contain AdjAx spinules (i.e., 74% of presynaptic boutons are SBBs, and 61% of these contain AdjAx spinules), and there are ∼30,000 synapses per pyramidal neuron (Megías et al., 2001), each CA1 pyramidal neuron would have ∼13,500 presynaptic boutons that envelop and putatively communicate (see *Potential Functions* below) with other axons and boutons. Furthermore, our observations of presynaptic inhibitory SBBs enveloping excitatory axons and spines (**Fig. 1G**_**5**_, **H**_**4**_ – **H**_**5**_), highlights the intriguing possibility that spinules may play a role in the rapid modulation of excitatory – inhibitory balance within CA1 microcircuits (Chevaleyre and Piskorowski, 2014).

### Synaptic Spinules Predict Synapse Strength

To our knowledge, only two other studies have examined the size of presynaptic boutons containing spinules, both presenting 2D evidence that SBBs are larger than adjacent non-SBBs (Erisir and Dreusicke, 2005; Campbell et al., 2020). Here, our 3D analyses revealed that SBB volumes and PSD surface areas are 2.5 and 1.6-times the size of non-SBBs, respectively, such that ∼50% of SBBs had volumes larger than 0.2 *μ*m^3^ and PSD areas larger than 0.12 *μ*m^2^, while only 4% of non-SBB volumes and 13% of their PSDs reached that size (**Fig. 3G** – **3J**). Accordingly, since presynaptic bouton volume and PSD surface area are directly related to synaptic strength and stability (Murthy et al., 2001; Ostroff et al., 2002; Branco et al., 2010; Holderith et al., 2012; Araya et al., 2014; Meyer et al., 2014; Quinn et al., 2019; Holler et al., 2021), we argue that SBBs represent the strongest and most stable excitatory presynaptic boutons within rat CA1.

### Putative Synaptic Spinule Functions

Surprisingly, we found that SBBs containing either small CT or large no-CT spinules were over 2-times larger than non-SBBs. This suggests two possibilities, either: (1) spinules aid in the stabilization and/or functional maturation of presynaptic boutons, or (2) spinules preferentially target and invaginate into already larger and stronger boutons. Regardless, the extreme differences in CT vs. no-CT spinule sizes and penetration strongly suggest they serve distinct functions. On average, large no-CT spinules project 450 nm into their SBBs (range: 90 nm – 1 *μ*m) and occupy 12% of their SBBs’ volume. Thus, the process of no-CT spinule envelopment (i.e., spinule projection and SBB invagination) involves a substantial level of actin cytoskeleton rearrangement and lipid bilayer modulation that together represent a large energy drain on the spinule-projecting and spinule-receiving neurons (Bernstein and Bamburg, 2003; Engl and Attwell, 2015). This level of energy and anatomical investment suggests a long-term relationship, with no-CT spinules potentially acting as structural anchors to enhance the stability of these non-canonical neuronal connections. Indeed, *in vitro*, the largest spinules are also the most stable (Zaccard et al., 2020). Additionally, SBBs share 2.4-times more parallel membrane interface with their large no-CT spinules than they do with their own PSDs (0.36 (spinule) vs. 0.15 (PSD) *μ*m^2^), making this interface a potential site for transmembrane communication (Wang et al., 2004). Alternatively, this large high-resistance membrane interface might allow for potent ephaptic communication between no-CT spinule projecting and receiving neurons, as large horizontal cell spinules do in vertebrate retina (Wagner and Djamgoz, 1993; Vroman et al., 2013). Finally, small CT spinules almost certainly transfer proteins and lipid bilayer between spinule and SBB, with the transendocytosis rate of these CT spinules increasing with high levels of neuronal activity (Sorra et al., 1998; Toni et al., 1999; Tao-Cheng et al., 2009).

## Notes

### Competing Interest Statement

The authors have declared no competing interest.

## References

Araya, R., Vogels, T.P., and Yuste, R. (2014). Activity-dependent dendritic spine neck changes are correlated with synaptic strength. Proc Natl Acad Sci U S A 111(28), E2895–2904. doi: 10.1073/pnas.1321869111.

Bernstein, B.W., and Bamburg, J.R. (2003). Actin-ATP Hydrolysis Is a Major Energy Drain for Neurons. The Journal of Neuroscience 23(1), 1–6. doi: 10.1523/jneurosci.23-01-00002.2003.

Branco, T., Marra, V., and Staras, K. (2010). Examining size-strength relationships at hippocampal synapses using an ultrastructural measurement of synaptic release probability. J Struct Biol 172(2), 203–210. doi: 10.1016/j.jsb.2009.10.014.

Calverley, R.K.S., and Jones, D.G. (1990). Contributions of dendritic spines and perforated synapses to synaptic plasticity. Brain Res Rev 15, 215–249.

Campbell, C., Lindhartsen, S., Knyaz, A., Erisir, A., and Nahmani, M. (2020). Cortical Presynaptic Boutons Progressively Engulf Spinules as They Mature. eNeuro 7(5). doi: 10.1523/ENEURO.0426-19.2020.

Case, N.M., Gray, E.G., and Young, J.Z. (1972). Ultrastructure and synaptic relations in the optic lobe of the brain of Eledone and Octopus. Journa of Ultrastructure Research 39(1), 115–123.

Chevaleyre, V., and Piskorowski, R. (2014). Modulating excitation through plasticity at inhibitory synapses. Frontiers in Cellular Neuroscience 8. doi: 10.3389/fncel.2014.00093.

Chirillo, M.A., Waters, M.S., Lindsey, L.F., Bourne, J.N., and Harris, K.M. (2019). Local resources of polyribosomes and SER promote synapse enlargement and spine clustering after long-term potentiation in adult rat hippocampus. Sci Rep 9(1), 3861. doi: 10.1038/s41598-019-40520-x.

Cover, K.K., and Mathur, B.N. (2021). Axo-axonic Synapses: Diversity in Neural Circuit Function. J Comp Neurol 529(9), 2391–2401.

Dall’Oglio, A., Dutra, A.C., Moreira, J.E., and Rasia-Filho, A.A. (2015). The human medial amygdala: structure, diversity, and complexity of dendritic spines. J Anat 227(4), 440–459. doi: 10.1111/joa.12358.

DeFelipe, J., and Fariñas, I. (1992). The pyramidal neuron of the cerebral cortex: Morphological and chemical characteristics of the synaptic inputs. Progress in Neurobiology 39(6), 563–607. doi: https://doi.org/10.1016/0301-0082(92)90015-7.

Engl, E., and Attwell, D. (2015). Non-signalling energy use in the brain. J Physiol 593(16), 3417–3429. doi: 10.1113/jphysiol.2014.282517.

Erisir, A., and Dreusicke, M. (2005). Quantitative morphology and postsynaptic targets of thalamocortical axons in critical period and adult ferret visual cortex. J Comp Neurol 485(1), 11–31.

Fiala, J.C. (2005). Reconstruct: a free editor for serial section microscopy. Journal of Microscopy 218, 52–61.

Geinisman, Y., deToledo-Morrell, L., and Morrell, F. (1994). Comparison of Structural Synaptic Modifications Induced by Long-Term Potentiation in the Hippocampal Dentate Gyrus of Young Adult and Aged Rats. Annals of the New York Academy of Sciences 747(1), 452–466.

Geinisman, Y., Detoledo-Morrell, L., Morrell, F., Heller, R.E., Rossi, M., and Parshall, R.F. (1993). Structural synaptic correlate of long-term potentiation: Formation of axospinous synapses with multiple, completely partitioned transmission zones. Hippocampus 3(4), 435–445. doi: 10.1002/hipo.450030405.

Geinisman, Y., Morrell, F., and deToledo-Morrell, L. (1987). Axospinous synapses with segmented postsynaptic densities: a morphologically distinct synaptic subtype contributing to the number of profiles of ‘perforated’ synapes visualized in random sections. Brain Res 423, 179–188.

Grillo, F.W., Neves, G., Walker, A., Vizcay-Barrena, G., Fleck, R.A., Branco, T., et al. (2018). A Distance-Dependent Distribution of Presynaptic Boutons Tunes Frequency-Dependent Dendritic Integration. Neuron 99(2), 275–282 e273. doi: 10.1016/j.neuron.2018.06.015.

Holderith, N., Lorincz, A., Katona, G., Rozsa, B., Kulik, A., Watanabe, M., et al. (2012). Release probability of hippocampal glutamatergic terminals scales with the size of the active zone. Nat Neurosci 15(7), 988–997. doi: 10.1038/nn.3137.

Holler, S., Kostinger, G., Martin, K.A.C., Schuhknecht, G.F.P., and Stratford, K.J. (2021). Structure and function of a neocortical synapse. Nature 591(7848), 111–116. doi: 10.1038/s41586-020-03134-2.

Inan, M., Blázquez-Llorca, L., Merchán-Pérez, A., Anderson, S.A., DeFelipe, J., and Yuste, R. (2013). Dense and Overlapping Innervation of Pyramidal Neurons by Chandelier Cells. The Journal of Neuroscience 33(5), 1907–1914. doi: 10.1523/jneurosci.4049-12.2013.

Knott, G., Rosset, S., and Cantoni, M. (2011). Focussed ion beam milling and scanning electron microscopy of brain tissue. J Vis Exp (53), e2588. doi: 10.3791/2588.

McCoy, J., and Nahmani, M. (2021). 3D Reconstruction and Analysis of Thin Subcellular Neuronal Structures using Focused-Ion Beam Scanning Electron Microscopy Data. J Vis Exp (175). doi: 10.3791/63030.

Megías, M., Emri, Z., Freund, T.F., and Gulyás, A.I. (2001). Total number and distribution of inhibitory and excitatory synapses on hippocampal CA1 pyramidal cells. Neuroscience 102(3), 527–540. doi: 10.1016/s0306-4522(00)00496-6.

Meyer, D., Bonhoeffer, T., and Scheuss, V. (2014). Balance and stability of synaptic structures during synaptic plasticity. Neuron 82(2), 430–443. doi: 10.1016/j.neuron.2014.02.031.

Murthy, V.N., Schikorski, T., Stevens, C.F., and Zhu, Y. (2001). Inactivity produces increases in neurotransmitter release and synapse size. Neuron 32, 673–682.

Ostroff, L.E., Fiala, J.C., Allwardt, B., and Harris, K.M. (2002). Polyribosomes redistribute from dendritic shafts into spines with enlarged synapses during LTP in developing rat hippocampal slices. Neuron 35(3), 535–545.

Perez-Alvarez, A., Yin, S., Schulze, C., Hammer, J.A., Wagner, W., and Oertner, T.G. (2020). Endoplasmic reticulum visits highly active spines and prevents runaway potentiation of synapses. Nat Commun 11(1), 5083. doi: 10.1038/s41467-020-18889-5.

Quinn, D.P., Kolar, A., Harris, S.A., Wigerius, M., Fawcett, J.P., and Krueger, S.R. (2019). The Stability of Glutamatergic Synapses Is Independent of Activity Level, but Predicted by Synapse Size. Front Cell Neurosci 13, 291. doi: 10.3389/fncel.2019.00291.

Schindelin, J., Arganda-Carreras, I., Frise, E., Kaynig, V., Longair, M., Pietzsch, T., et al. (2012). Fiji: an open-source platform for biological-image analysis. Nature Methods 9(7), 676–682. doi: 10.1038/nmeth.2019.

Schuster, T., Krug, M., and Wenzel, J. (1990). Spinules in axospinous synapses of the rat dentate gyrus: changes in density following long-term potentiation. Brain Research 523(1), 171–174. doi: https://doi.org/10.1016/0006-8993(90)91654-Y.

Shepherd, G.M., and Harris, K.M. (1998). Three-Dimensional Structure and Composition of CA3 to CA1 Axons in Rat Hippocampal Slices: Implications for Presynaptic Connectivity and Compartmentalization. J Neurosci 18(20), 8300–8310.

Somogyi, P., Freund, T.F., and Cowey, A. (1982). The axo-axonic interneuron in the cerebral cortex of the rat, cat and monkey. Neuroscience 7(11), 2577–2607. doi: 10.1016/0306-4522(82)90086-0.

Sorra, K.E., Fiala, J.C., and Harris, K.M. (1998). Critical assessment of the involvement of perforations, spinules, and spine branching in hippocampal synpase formation. J Comp Neurol 398, 225–240.

Spacek, J., and Harris, K.M. (2004). Trans-endocytosis via spinules in adult rat hippocampus. J Neurosci 24(17), 4233–4241. doi: 10.1523/JNEUROSCI.0287-04.2004.

Tao-Cheng, J.H., Dosemeci, A., Gallant, P.E., Miller, S., Galbraith, J.A., Winters, C.A., et al. (2009). Rapid turnover of spinules at synaptic terminals. Neuroscience 160(1), 42–50. doi: 10.1016/j.neuroscience.2009.02.031.

Tarrant, S.B., and Routtenberg, A. (1977). The synaptic spinule in the dendritic spine: electron microscopic study of the hippocampal dentate gyrus. Tissue Cell 9(3), 461–473.

Toni, N., Buchs, P.A., Nikonenko, I., Bron, C.R., and Muller, D. (1999). LTP promotes formation of multiple spine synapses between a single axon terminal and a dendrite. Nature 402(6760), 421–425.

Vroman, R., Klaassen, L.J., and Kamermans, M. (2013). Ephaptic communication in the vertebrate retina. Front Hum Neurosci 7, 612. doi: 10.3389/fnhum.2013.00612.

Wagner, H., and Djamgoz, M. (1993). Spinules: a case for retinal synaptic plasticity. Trends Neurosci 16(6), 201–206.

Wang, Y., Chan, S.L., Miele, L., Yao, P.J., Mackes, J., Ingram, D.K., et al. (2004). Involvement of Notch signaling in hippocampal synaptic plasticity. Proceedings of the National Academy of Sciences 101(25), 9458–9462. doi: doi:10.1073/pnas.0308126101.

Westfall, J.A. (1970). Ultrastructure of synapses in a primitive coelenterate. Journal of ultrastructure research 32(3), 237–246. doi: 10.1016/s0022-5320(70)80004-1.

Westrum, L.E., and Blackstad, T.W. (1962). An electron microscopic study of the stratum radiatum of the rat hippocampus (regio superior, CA 1) with particular emphasis on synaptology. J Comp Neurol 119, 281–309.

Zaccard, C.R., Shapiro, L., Martin-de-Saavedra, M.D., Pratt, C., Myczek, K., Song, A., et al. (2020). Rapid 3D Enhanced Resolution Microscopy Reveals Diversity in Dendritic Spinule Dynamics, Regulation, and Function. Neuron. doi: 10.1016/j.neuron.2020.04.025.

Zhang, N., and Houser, C.R. (1999). Ultrastructural Localization of Dynorphin in the Dentate Gyrus in Human Temporal Lope Epilepsy: A Study of Reorganized Mossy Fiber Synapses. J Comp Neurol 405, 472–490.

